# Global DNA methylation, as estimated in blood samples, does not correlate with variations of body condition, anatomical features and hematological parameters in American bullfrogs *(Lithobates catesbeianus)* kept captive under distinct environmental conditions

**DOI:** 10.1101/156265

**Authors:** Braulio Ayala García, Alma L. Fuentes-Farías, Gabriel Gutiérrez Ospina

## Abstract

Different levels of Global DNA Methylation (GDM) could have facilitated the emergence of new species, without relying on gene mutations, through promoting ontogenetic phenotypic plasticity. If this assertion was correct, one could expect individuals of the same species living under distinct environmental conditions to be genetically similar, but having different GDM levels and being phenotypically divergent. We tested this presumption by studying the relationship between variability of functional morphological traits and GDM levels in American bullfrog *(Lithobates catesbeianus)*, in green houses located in two geographical sites. Our analyses revealed that body linear morphometry, skull geometry, scaled mass index, packed cell volume and neutrophil counts differed significantly among males and females within and between localities. GDM, nonetheless, was rather similar among sex and locality groups. These results show that levels of GDM, at least under our experimental contexts, does not correlate with functional morphological trait variability.

## Introduction

Shifts in DNA methylation are one of the epigenetic processes that are presumed to contribute to generate non-mutational phenotype variability, within and across animal populations by coding environmental information(Baerwald *et al.*, 2016). It is by modulating the degree of transcriptional elongation(Rountree and Selker, 1997) and chromatin compaction/relaxation (Bogdanovic *et al.*, 2011), that DNA methylation regulate patterns of gene expression and ontogeny (Dunican *et al.*, 2008). Methylation takes place in the fifth carbon of cytosine residues (i.e., 5-methyl-cytosine), within gene bodies, enhancers and promoter regions, in intergenic regions, among others (Stancheva *et al.*, 2002). The chemical reaction involved in this process is catalyzed by a family of methyl-transferase enzymes(Stancheva, Hensey and Meehan, 2001).

Different levels of Global DNA methylation (GDM) occur among vertebrate classes and orders(Vanyushin *et al.*, 1973). Nowadays, it is presumed that shifts in global DNA methylation (GDM) are important for understanding phenotype plasticity promoted by daily live interaction of the organism with the environment(Zhu *et al.*, 2012), and hence, favor non-mutational phenotype variability during vertebrate evolution(Ponger and Li, 2005). For instance, monotremes have the highest levels of methylated DNA, followed by placentals and marsupials. In a similar vein, amphibian and fish DNA is about twice as methylated as that of reptiles, birds and mammals(Jabbari *et al.*, 1997). Therefore, shifts in GDM are thouhght to have contributed in driving the “cold to warm transition” during vertebrate evolution(Varriale and Bernardi, 2006b). That environmental factors may shift GDM, is supported by reports that show that fish living near the poles have higher values of GDM, as estimated in blood samples, than those reported for fish living in temperate or tropical areas(Varriale and Bernardi, 2006a). Since differences in GDM were reported to occur in fish species whose phylogenetic relatedness is high, the results reported further support that interspecies GDM in part reflects differential environmental exposure(Varriale, 2014). However, it remains unclear whether this might be true for phenotype variability observed in individuals of a single species that are exposed to distinct environments.

Domestication and artificial selection has traditional been a powerful tool to infer rules governing the evolution of species(Rivas Sanchez and Rivas Sanchez, 2015). Here, we turn to this strategy to evaluate whether trait variability, as analized through linear morphometry, skull geometric morphometry, body condition, whole white blood cell count, packed cell volume (PCV) and differential leukocyte counting, is correlated with GDM levels in bullfrogs subjected to distinct environments. Bullfrogs are an adequate experimental unit since their colonies display reduced genetic diversity(Bai *et al.*, 2012), but a great deal of phenotypic plasticity. In fact, this ability has made bullfrogs rather successful dwellers, They have literally colonized a variety of habitats throughout the globe (Govindarajulu, Price and Anholt, 2006).

## Methods

### Animals

A total of sixty bullfrogs *(L. catesbeianus;* 1 year old; 15 males / 15 females per locality, n=15) were obtained from “La Purísima” (LPu) and “San Pedro Tlaltizapán” (SPt), two rustic greenhouses located at the State of Michoacán (19°52’11“N, 101°01’23“W) and at the Estado de México (19°20’11“N, 99°49’79“W), respectively. LPu (6033,465 ft) is about 6°C warmer than in SPt (8448,163 ft) through the year. In both sites male and female frogs were kept captive in groups of 100 individuals that coinhabited large, cement-made tanks (2.5m^2^). Frogs were fed twice a day with commercial trout food provided by a local supplier (El Pedregal, Toluca, Estado de México; composition: 45% protein, 16% fat, 2.5% fiber, 12% ash, 12% humidity). Frogs were transferred to the laboratory in 200 liters plastic containers. Transportation took no more than six hours from frog capturing to freeing them in the laboratory enclosure. Frogs were kept under laboratory condition during 12 hours before sacrifice. All efforts were made to minimize stress; for instance, frogs were kept in a quiet and dark room and were not disturbed until sacrifice. Animal handling and experimentation followed federal guidelines recommended by the Mexican Official Norm on production, housing and handling of laboratory animals (NOM-62-ZOO-1999). All sections of this paper were made according to the “ARRIVE guidelines for reporting animal research” (Kilkenny *et al.*, 2010).

### Assessing body condition

Body condition was estimated in LPu and SPt, male and female bullfrogs by using the Scaled Mass Index(MacCracken and Stebbings, 2012). This condition index (CI), is better than the OLS residual method and other CI’s because it correlates better with fat reserves and lean dry mass in reptiles and mammals (Peig, Green and Ame, 2009). The SMI was calculated using the following formula:

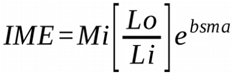

Where *Mi* and *Li* stand for individual mass and snout-vent length, *Lo* is the arithmetic mean of all the animals sampled, and *“bsma”* is the coefficient of a standard major axis regression for all the mass and length data from both male and female, mature and immature individuals.

### Hematological parameters

Blood cell composition was evaluated in blood samples obtained from juvenile, male and female bullfrogs anesthetized with ice following their decapitation. Blood was collected in EDTA-treated tubes and stored at -4°C. Packed cell volume (PCV) was estimated after centrifuging 2ml of blood at 4,500 rpm for 15 minutes, and then measuring the volume of both serum and hematocrit. Leukocyte counts (WBC) were done in a hemocytometer after conventional Turk-solution treatment (for at least 5 minutes) of blood samples. Leukocyte differential counts were done on blood smears stained with Wright reagent for 5 minutes. A total of 100 cells in each smear were counted and classified as lymphocytes, neutrophils, monocytes, basophils and eosinophils, following specific guidelines of amphibian hematology (Allender and Fry, 2008).

### Morphometries

Linear morphometric analysis was done on 7 lineal measurements taken with a precision Veneer caliper (0.01 cm error). Measures include snout-vent length, head width and length, timpanum and eye diameter, leg size (femur plus tibio-fibula), feet (tarus plus metatarsus). For the geometric analysis, we used 10 skulls (n=10) that were previously cleaned with dissecting tools and using commercially bleacher for a few minutes to avoid damage. Digital photographies from the ventral aspect of the skull were taken and used to place ten landmarks as suggested for the *Rana* genus (Fig. S1). The positioning of the landmarks was done according to (Larson, 2002, 2005). TpsUtil and Tps2 (James Rohlf, Stony Brook University) software was used to generate landmark information from the digital images. Subsequently, landmark configurations for all individuals were superimposed and aligned acording to the ordinary least-squares Procrustes method, whit which Procrustes and Mahalanobis distance between and within groups was calculated.

### Estimating global DNA methylation (GDM)

DNA was isolated from blood samples using the propanol extraction protocol (Miller, Dykes and Polesky, 1988). DNA concentration and purity was estimated using a Nano-drop 1000 (Thermo-Fisher, Waltham MA, USA). DNA integrity was evaluated using 1% agarose gel electrophoresis. The percentage of 5-methylcitosine was estimated using an ELISA kit following manufacturer’s protocol (Zymo D5326, Irving CA, USA). Briefly, 100ng of double stranded DNA were denatured for 5 minutes in a thermocycler (Axygen Union City CA, USA). Then, DNA samples were adsorbed each to the walls of wells of 96-well plates. Each assays was conducted by duplicate. After blocking, anti-5-mC primary monoclonal antibody (1:2,000) and HRP secondary antibody (1:1,000) were mixed, and wells were incubated with 100 µl of this solution for 1 hour at 37°C (information about animals used to raise. Following several washes, HRP activity was revealed by adding 100 µl of manufacturer’s HRP developer during 10 to 60 minutes at room temperature. Absorbance was estimated at 480 nm with the aid of an ELISA plaque reader (Biotek, Winooski VT, USA). The percentage of DNA methylation in each sample was calculated after fitting logarithmic second-order regression on the manufacturer’s standard curve absorbance values.

### Statistical analysis and software

Morphological data from both groups were compared by using a multivariate analysis conducted through canonical discriminant tests (CDA). We used JMP statistical software (SAS, version 10) and MorphoJ 2.0 (Obtained from Klingenberg’s lab official site:http://www.flywings.org.uk/morphoj_page.htm). This method was used instead of principal component analysis (PCA) because it maximizes the degree to which pre-defined groups can be distinguished. To find any difference between experimental groups on linear morphometrics, Wilk’s Lambda (λ) was calculated at p< 0.05. For geometric analysis, Mahalanobis distance was calculated after ordinary Procsrustes alignment (MDA. MorphoJ 2.0). In both cases, canonical component 1 and 2 were also used to conduct a 2-way ANOVA, to test for possible sex and locality effects on morphometric variables. Body condition, whole white blood cell count, packed cell volume and differential leukocyte number are presented as mean ± s.e.m. After testing equal variance, Shapiro-Wilk test was used to ensure normality of the data. To test differences between groups from body condition, hematology and GDM variables, we used a One-way ANOVA followed by a Tuckey post-hoc test, with an alpha level of < 0.05. R 3.1.1 was used for statistical analysis and Sigma Plot 12 to create graphs.

## Results

In this experiments, we tested the possible correlation between phenotypic variability and GDM by using bullfrogs from two localities, a species that display a great deal of environmentally driven phenotypic plasticity in the absence of genetic mutations. Both female and male bullfrogs were larger in SPt than in LPu. The overall morphometry between females from both sites was similar, whereas differences could be seen in their male counterparts (λ =0.139; F 3.68, p<0.001; Fig. 1A). Differences between female and male frogs from LPu were greater than in Spt, thus sex and locality exert a combined effect. (2-ANOVA; p=0.0004). This results suggests that sexual dimorphism is present at both sites but is more noticeable in LPu. On the other hand, skull geometry behaved differently in both populations (Fig. 1B). In LPu, there was a sharp difference between female and male skull shape (p<0.0001). This was not observed in SPt (MDA; p=0.1). Lastly, even though no differences in skull shape were documented between females of both localities (MDA; p=0.2), males did differ significantly (MDA; p<0.0001).

**Figure 1.**
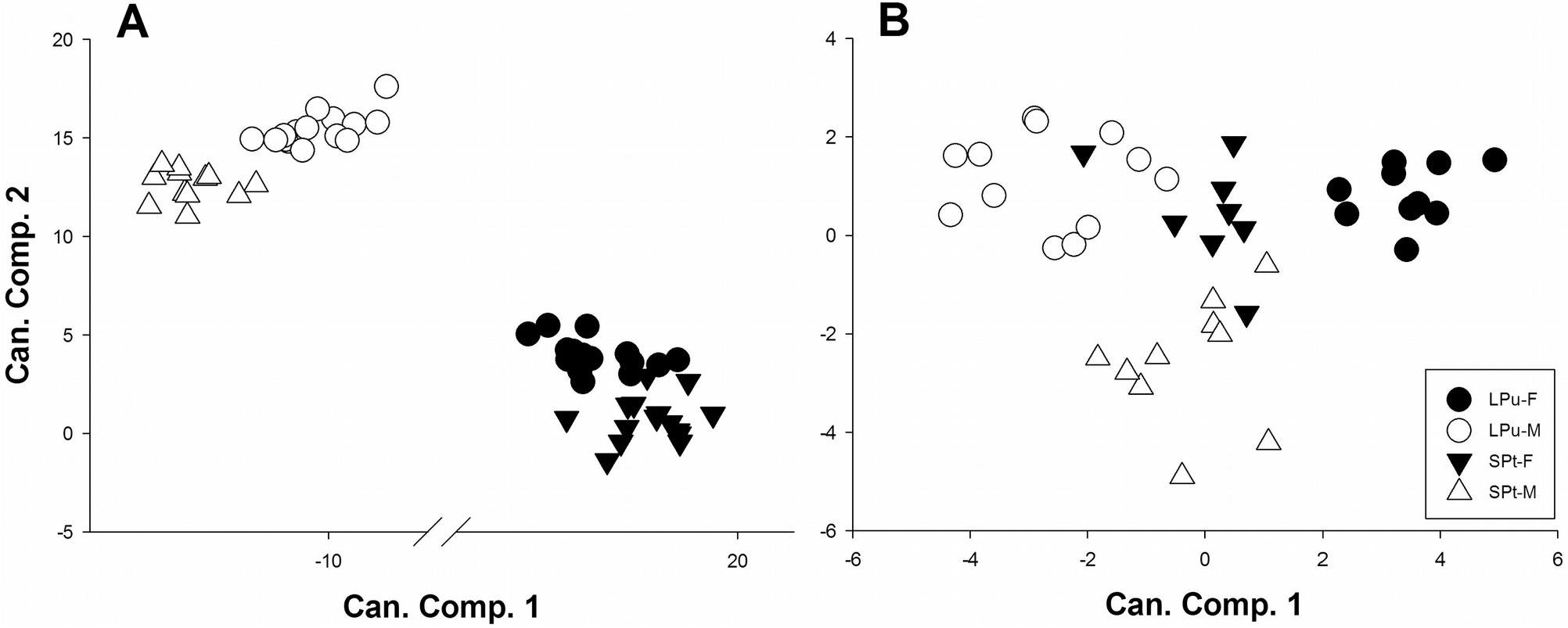
Linear and geometric morphometrics. Canonical variate analysis was performed on 7 lineal measurements for linear morphometrics (A), and on ten landmark extracted from ventral digital photos of the skull (B) of female and male bullfrogs. Canonical component (CP) 1 was plotted against CP2 for every frog. We used 30 frogs (15 female and 15 male, n=15) for linear morphometrics and 20 skulls ( 10 female, and 10 male, n=10) from each of the two localities. Black circles correspond to LPu females, open circles to LPu males, black triangles to SPt females and open triangles to males (n = 15).

Body condition, as estimated by the Scaled Mass Index, differed between LPu and SPt bullfrogs (Table 1). Both male and female frogs from SPt had a higher body condition than LPu ones (ANOVA, p<0.0001). As shown in Table 1, PCV levels were higher in both male and female frogs from LPu than SPt animals (ANOVA; p<0.0001). WBC counts on the contrary, did not differ between among groups (ANOVA; p=0.96). Overall, neutrophil number was greater in LPu than in SPt. In the former locality, nonetheless, neutrophil number was higher in male than in female frogs (ANOVA; p<0.05. Table 2). In SPt, monocyte number was 3.4-fold higher in males than in females. In addition, monocyte number of males from SPt was 6.9-fold higher than that observed in specimens of both sexes raised in LPu (Table 2).

**Table 1.**
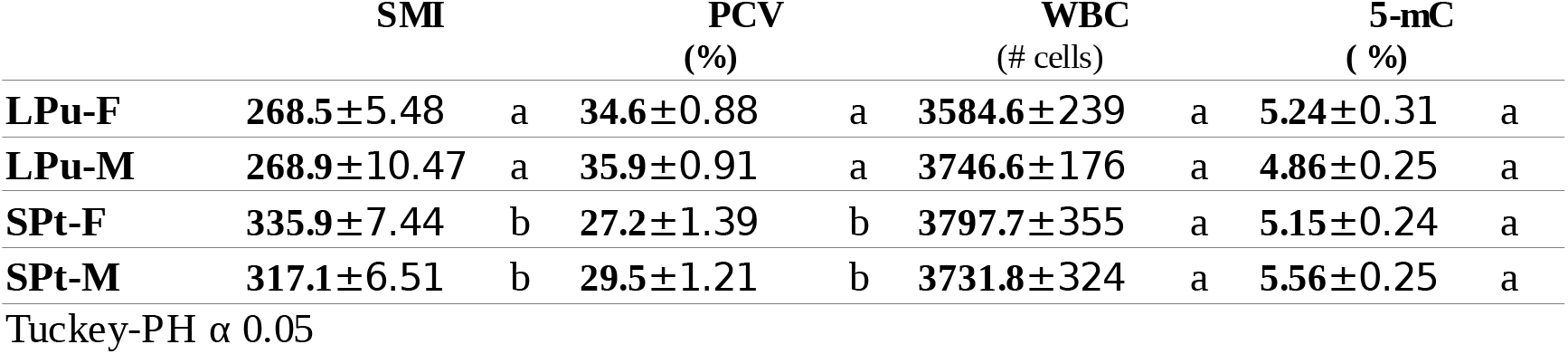
Physiological parameters and global content of 5-mC. Scaled mass index, packed cell volume, whole blood cell count and the percentage of 5-mC were measured in 15 male and 15 femal frogs from two localities (LPu and Spt, n=15). Mean and s.e.m. are reported. Letters indicate statistical significance after Tuckey post-hoc test on a One way-ANOVA. For PCV, WBC and 5-mC, blood samples were used (for sample processing see materials and methods).

**Table 2.** Differential leukocyte count. The five principal cell types in blood samples were measured in 15 male and 15 femal frogs from two localities using a conventional Wright staining technique (LPu and SPt, n=15). Mean and s.e.m. are reported, and letters indicate statistical significance after Tuckey post-hoc test on a One way-ANOVA.

|  | Neutrophils |  | Lymphocytes |  | Monocytes |  | Basophils |  | Eosinophils |  |
| --- | --- | --- | --- | --- | --- | --- | --- | --- | --- | --- |
| **LPu-F** | 1154.27±169.9 | ab | 1997.16±225.7 | a | 36.61±8.63 | a | 230.55±51.1 | a | 125.83±26.16 | a |
| **LPu-M** | 1464.53±128.1 | b | 1791.14±103.7 | a | 31.07±7.6 | a | 233.17±49 | a | 197.92±49 | a |
| **Spt-F** | 730.84±63.5 | a | 2093.4±236.3 | a | 68.69±19.8 | a | 413.32±75.9 | a | 191±29.4 | a |
| **Spt-M** | 933.61±120 | a | 1927.84±189.7 | a | 233.65±40.8 | b | 349.13±53 | a | 236.27±50 | a |

Tuckey-PH $\alpha$ 0.05

Global DNA methylation estimated in blood samples showed neither differences between sex or locality (Table 1). Moreover, there was no correlation between this variable with any of the others. A multivariate analysis of the phenotypic variation in response of GDM did not any significant correlation (not shown)

### Discussion

Shifts in GDM are thought to contribute in generating phenotype variability during evolutionary processes (Jabbari *et al.*, 1997; Varriale and Bernardi, 2006b). Even though significant differences in GDM seem to associate with trait variability when comparing animal classes (Jabbari *et al.*, 1997), this presumption may not be valid for explaining trait variability among individuals belonging to the same species. Accordingly, here we showed that bullfrogs raised in two localities differ significantly in body linear morphometry, skull geometry (Fig. 1), scaled mass index, packed cell volume and neutrophil counts (Tables 1 - 2). GDM, however, was similar between all groups, and not correlated with other variables (Table 1). Hence GDM, under current experimental context, does not correlate with trait variability.

The fact that GDM did not correlate with any phenotypic variable is at odds with previous studies. For instance, fish species belonging to the same family are known to have different GDM levels depending on the temperature they are exposed to. This indeed is the case for *Symphodus tinca*, a Mediterranean fish that exhibits the highest levels of GDM among the members of Labridae family (Varriale and Bernardi, 2006a). Moreover, expecting that GDM levels could differ between bullfrogs from SPt and LPu was a reasonable presumption since even different organ within single individuals may have different GDM levels, as shown in rats, mice and monkeys (Gama-Sosa *et al.*, 1983). Although we do not currently have an explanation for these counterintuitive results, it is fair to say that we cannot rule out fully that global methylation is unrelated with trait variability since differential methylation may occur in distinct genes throughout the genome for individuals coming from one locality or the other. Accordingly, ELISA GDM analysis may not be the correct tool for approaching inter-individual variation within single species. It may be more instructive to analyze global patterns of DNA methylation as shown for fish (Baerwald *et al.*, 2016), birds (Liebl *et al.*, 2013), reptiles (Venegas *et al.*, 2016) and mammals (Chang *et al.*, 2006).

Recent evidence suggests that GDM may participate in the process of generating non-mutational phenotypic variability in natural conditions (Varriale, 2014). Bullfrogs living out of their native range exhibit a highly variable phenotype that could be attributed to the effect of being exposed to distinct environmental niches (Govindarajulu, Price and Anholt, 2006). In this work, we found no correlation between morphological nor physiological traits with levels of GDM. However, our results show that intraspecific sex variability is also present in bullfrogs even under relatively controlled conditions (Figs. 1A-B), thus suggesting that sexual dimorphism in bullfrogs can be influenced by environmental factors. Future studies must establish whether global patterns of DNA methylation can explain these interesting findings.

## Acknowledgements

We thank Dr. Jesús Ramírez Santos and Dr. Margarita Gómez Chavarín for their technical support. We also thank Posgrado en Ciencias de la Producción y de la Salud Animal (UNAM), and the Consejo Nacional de Ciencia y Tecnología, through which this study was conducted.

## Competing interests

The authors declare no competing financial interests regarding the elaboration of the study and the preparation of the manuscript.

## Author contribution

First and last author contributed equally to the presented study. Second author contributed with preparation of the manuscript and data analysis.

## Funding

This study was funded with institutional resources if the Instituto de Investigaciones Biomédicas of the Universidad Nacional Autónoma de México. The first author was a PhD student granted by the Consejo Nacional de Ciencia de Ciencia y Tecnología (CONACYT).

## Data availability

The data used for the analysis of this study can be found in Dryad: (after submission

## References

Allender, M. C. and Fry, M. M. (2008) Amphibian hematology. The veterinary clinics of North America. Exotic animal practice. 11(3), 463–80. doi: 10.1016/j.cvex.2008.03.006.

Baerwald, M. R., Meek, M. H., Stephens, M. R., Nagarajan, R. P., Goodbla, A. M., Tomalty, K. M. H., Thorgaard, G. H., May, B. and Nichols, K. M. (2016). Migration-related phenotypic divergence is associated with epigenetic modifications in rainbow trout, *Molecular Ecology*. 25(8), 1785–1800. doi: 10.1111/mec.13231.

Bai, C., Ke, Z., Consuegra, S., Liu, X. and Li, Y. (2012). The role of founder effects on the genetic structure of the invasive bullfrog *(Lithobates catesbeianaus)* in China. Biological Invasions, 14(9), 1785–1796. doi: 10.1007/s10530-012-0189-x.

Bogdanovic, O., Long, S. W., van Heeringen, S. J., Brinkman, A. B., Gómez-Skarmeta, J. L., Stunnenberg, H. G., Jones, P. L. and Veenstra, G. J. C. (2011). Temporal uncoupling of the DNA methylome and transcriptional repression during embryogenesis. Genome research, 21, 1313–1327. doi: 10.1101/gr.114843.110.

Chang, H.-S., Anway, M. D., Rekow, S. S. and Skinner, M. K. (2006). Transgenerational epigenetic imprinting of the male germline by endocrine disruptor exposure during gonadal sex determination. Endocrinology. 147(12),5524–41. doi: 10.1210/en.2006-0987.

Dunican, D. S., Ruzov, A., Hackett, J. a and Meehan, R. R. (2008). xDnmt1 regulates transcriptional silencing in pre-MBT Xenopus embryos independently of its catalytic function. Development (Cambridge, England). 135(7), 295–302. doi: 10.1242/dev.016402.

Gama-Sosa, M. A., Midgett, R. M., Slagel, V. A., Githens, S., Kuo, K. C., Gehrke, C. W. and Ehrlich, M. (1983). Tissue-specific differences in DNA methylation in various mammals. BBA - Gene Structure and Expression. 740(2), 212–219. doi: 10.1016/0167-4781(83)90079-9.

Govindarajulu, P., Price, W. M. S. and Anholt, B. R. (2006). Introduced Bullfrogs *(Rana catesbeiana)* in Western Canada: Has Their Ecology Diverged?. Journal of Herpetology. 40(2), 249–260. doi: 10.1670/68-05A.1.

Jabbari, K., Cacciò, S., Païs De Barros, J. P., Desgrès, J. and Bernardi, G. (1997). Evolutionary changes in CpG and methylation levels in the genome of vertebrates. Gene,. 205(1-2), 109–118. doi: 10.1016/S0378-1119(97)00475-7.

Kilkenny, C., Browne, W. J., Cuthill, I. C., Emerson, M. and Altman, D. G. (2010). Improving Bioscience Research Reporting: The ARRIVE Guidelines for Reporting Animal Research. PLoS Biology. 8(6), e1000412. doi: 10.1371/journal.pbio.1000412.

Larson, P. M. (2002). Chondrocranial development in larval *Rana sylvatica* (Anura: Ranidae): Morphometric analysis of cranial allometry and ontogenetic shape change. Journal of Morphology. 252(2), 131–144. doi: 10.1002/jmor.1095.

Larson, P. M. (2005). Ontogeny, phylogeny, and morphology in anuran larvae: morphometric analysis of cranial development and evolution in *Rana* tadpoles (Anura: Ranidae). Journal of morphology. 264(1), 34–52. doi: 10.1002/jmor.10313.

Liebl, A. L., Schrey, A. W., Richards, C. L. and Martin, L. B. (2013). Patterns of DNA Methylation Throughout a Range Expansion of an Introduced Songbird. Integrative and comparative biology. 53(2),351–358. doi: 10.1093/icb/ict007.

MacCracken, J. G. and Stebbings, J. L. (2012). Test of a Body Condition Index with Amphibians. Journal of Herpetology. 46(3), 346–350. doi: 10.1670/10-292.

Miller, S. A., Dykes, D. D. and Polesky, H. F. (1988). A simple salting out procedure for extracting DNA from human nucleated cells. Nucleic Acids Research. 16(3), 1215–1215. doi: 10.1093/nar/16.3.1215.

Peig, J., Green, A. J. and Ame, C. (2009). New perspectives for estimating body condition from mass / length data: the scaled mass index as an alternative method. 118(12), 1883–1891. doi: 10.1111/j.1600-0706.2009.17643.x.

Ponger, L. and Li, W.-H. (2005). Evolutionary diversification of DNA methyltransferases in eukaryotic genomes. Molecular biology and evolution. 22(4), 1119–28. doi: 10.1093/molbev/msi098.

Rivas Sanchez, D. F. and Rivas Sanchez, D. F. (2015) ‘Identifying potential for fisheries-induced evolution on behavioral traits of a Skagerrak cod *(Gadus morhua)* population. Master thesis, University of Oslo, 2015. Available at: https://www.duo.uio.no/handle/10852/46128 (Accessed: 13 June 2017).

Rountree, M. R. and Selker, E. U. (1997). DNA methylation inhibits elongation but not initiation of transcription in Neurospora crassa. Genes & development. 11(18), 2383-95.

Stancheva, I., El-Maarri, O., Walter, J., Niveleau, A. and Meehan, R. R. (2002) .DNA methylation at promoter regions regulates the timing of gene activation in *Xenopus laevis* embryos. Developmental biology. 243(1), 155–65. doi: 10.1006/dbio.2001.0560.

Stancheva, I., Hensey, C. and Meehan, R. R. (2001). Loss of the maintenance methyltransferase, xDnmt1, induces apoptosis in *Xenopus* embryos. 20(8), 1963–1973. doi: 10.1093/emboj/20.8.196.

Vanyushin, B. F., Mazin, A. L., Vasilyev, V. K. and Belozersky, A. N. (1973) .The content of 5- methylcytosine in animal DNA: The species and tissue specificity. BBA Section Nucleic Acids And Protein Synthesis. 299(3), 397–403. doi: 10.1016/0005-2787(73)90264-5.

Varriale, A. (2014). DNA Methylation, Epigenetics, and Evolution in Vertebrates: Facts and Challenges. International journal of evolutionary biology. 2014, 1–7. doi: 10.1155/2014/475981.

Varriale, A. and Bernardi, G. (2006a). DNA methylation and body temperature in fishes. Gene. 385, 111–121. doi: 10.1016/j.gene.2006.05.031.

Varriale, A. and Bernardi, G. (2006b). DNA methylation in reptiles. Gene. 385, 122–127. doi: 10.1016/j.gene.2006.05.034.

Venegas, D., Marmolejo-Valencia, A., Valdes-Quezada, C., Govenzensky, T., Recillas-Targa, F. and Merchant-Larios, H. (2016). Dimorphic DNA methylation during temperature-dependent sex determination in the sea turtle *Lepidochelys olivacea*. General and Comparative Endocrinology. 236, 35–41. doi: 10.1016/j.ygcen.2016.06.026.

Zhu, Z. Z., Hou, L., Bollati, V., Tarantini, L., Marinelli, B., Cantone, L., Yang, A. S., Vokonas, P., Lissowska, J., Fustinoni, S., Pesatori, A. C., Bonzini, M., Apostoli, P., Costa, G., Bertazzi, P. A., Chow, W. H., Schwartz, J. and Baccarelli, A. (2012). Predictors of global methylation levels in blood DNA of healthy subjects: A combined analysis. International Journal of Epidemiology. 41(1), 126–139. doi: 10.1093/ije/dyq154.

